# Automated map sharpening by maximization of detail and connectivity

**DOI:** 10.1101/247049

**Authors:** Thomas C. Terwilliger, Oleg Sobolev, Pavel V. Afonine, Paul D. Adams

## Abstract

**Synopsis:** A procedure for optimizing the sharpening of a map based on maximizing the level of detail and connectivity of the map is developed and applied to 361 pairs of deposited cryo-EM maps and associated models.

**Abstract:** We present an algorithm for automatic map sharpening that is based on optimization of detail and connectivity of the sharpened map. The detail in the map is reflected in the surface area of an iso-contour surface that contains a fixed fraction of the volume of the map, where a map with high level of detail has a high surface area. The connectivity of the sharpened map is reflected in the number of connected regions defined by the same iso-contour surfaces, where a map with high connectivity has a small number of connected regions. By combining these two measures in a metric we term “adjusted surface area”, we can evaluate map quality in an automated fashion. We use this metric to choose optimal map sharpening parameters without reference to a model or other interpretations of the map. Map sharpening by optimization of adjusted surface area can be carried out for a map as a whole or it can be carried out locally, yielding a locally-sharpened map. To evaluate the performance of various approaches, we use a simple metric based on map-model correlation that can reproduce visual choices of optimally-sharpened maps. The map-model correlation is calculated using a model with B-factors (atomic displacement factors, ADP) set to zero. We use this model-based metric to evaluate map sharpening, use it to evaluate map sharpening approaches and find that optimization of adjusted surface area can be an effective tool for map sharpening.

## 1. Introduction

Current methods for single-particle reconstruction of cryo-EM maps are capable of yielding maps with resolution often better than 4.5 Å and sometimes as high as 2 Å or better (c.f., Kuhlbrandt, 2014; Baldwin et al., 2018). The level of noise in the images used in reconstruction and in the resulting maps is highly resolution-dependent. For this reason it is standard practice to represent a map as a Fourier series, estimate the signal and noise in the reconstruction as a function of resolution, and use this information to weight the Fourier terms as a function of resolution to maximize the interpretability of the final sharpened (or blurred) map (Rosenthal and Henderson, 2003). As the various errors in reconstruction are difficult to estimate accurately, other approaches for resolution-dependent weighting have been considered as well. For example, the *Phenix* feature-enhanced maps (*FEM,* Afonine et al., 2015) and the model-building software *Coot* (Emsley et al., 2010) use maximization of the kurtosis of a map to choose an overall sharpening *B*_*sharpen*_ (an exponential factor applied to Fourier terms, cf. DeLaBarre and Brunger, 2006; Wlodawer et al., 2008). Nicholls et al. (2012) develop procedures for optimizing anisotropic versions of displacement factors based on considering sharpening as an inverse deblurring problem. Joseph et al. (2017) use the method of Rosenthal and Henderson for map sharpening during the refinement of macromolecular structures. Burnley et al. (2017) have recently noted that the challenge of optimizing map sharpening is an open one with the comment that, “Presently, the optimum sharpening coefficient (where ‘optimum’ means maximizing the interpretable density features) cannot be analytically determined either locally or globally, although attempts are ongoing.” Model-based approaches have been used for this purpose, however. Falke (2005) used the resolution-dependence of a model-based map in a sharpening procedure and recently Jakobi et al. (2017) applied such a sharpening procedure locally to optimize the contrast and interpretability of density maps. Sharpening is also commonly applied in X-ray crystal structure analysis. DeLaBarre and Brunger (2006) suggested strongly sharpening low-resolution maps; their procedure leads to an overall B-value *B*_*iso*_ (isotropic Wilson B-factor, iso stands for isotropic here; closely related to B-factors or atomic displacement factors, cf. DeLaBarre and Brunger, 2006; Wlodawer et al., 2008) of about zero. The *Phenix* (Adams et al., 2010) tools *autosol* (Terwilliger et al., 2009) and *autobuild* (Terwilliger et al., 2010) use map sharpening routinely in automated map interpretation and sharpened maps to an overall B-value numerically given by 10 times the resolution in Å units (e.g., *B*_*iso*_ = 40 Å^2^ at a resolution of 4 Å; Terwilliger et al., 2008). Liu and Xiong (2014) applied sharpening to nearly 2000 X-ray maps and found that map-model correlation could generally be improved through map sharpening.

Here we present a model-free algorithm for optimizing the sharpening of a map that is based on simultaneously maximizing the level of detail in the map and the connectivity of the map. We show that this procedure can be an effective tool for map sharpening.

## 2. Methods

### 2.1. Map sharpening and blurring

Prior to map sharpening or blurring, maps are (by default) corrected for anisotropy using the *Phenix* tool *phenix.xtriage* (Zwart et al., 2005), where the final isotropic B-value *B*_*iso*_ is set to be the average of the 3 diagonal terms in the matrix representing the anisotropic B-factor.

We use a four-parameter function for map sharpening. The map is represented as a Fourier series and is typically sharpened at lower resolution and then blurred at higher resolution though the map can be blurred at all resolutions as well. The four parameters are: a sharpening B-factor (*B*_*sharpen*_) applied at lower resolutions, a blurring B-factor (*B*_*blur*_) applied at high resolution, a transition resolution (*d*_*cut*_) and a transition parameter (*k*) used to define over what resolution range the transition between sharpening and blurring occurs. Note that by “sharpening” here we mean either overall sharpening or blurring. If the map is sharpened the sharpening B-factor (*B*_*sharpen*_) is positive, if the map is blurred it is negative. If the map is blurred, no additional blurring B-factor is applied at high resolution.

The sharpening B-factor (*B*_*sharpen*_) is applied to amplitudes in the Fourier series representing the map through a resolution-dependent sharpening scale factor *A*_*sharpen*_ *(d)*, where *d* is the resolution of a Fourier term:

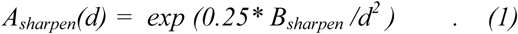

If the value of the B-factor (*B*_*sharpen*_) is positive so that the map is being sharpened (amplitudes increasing at high resolution), a blurring scale factor is applied at high resolution as a kind of soft resolution limit. The blurring scale factor *A*_*blur*_*(d)* is given by,

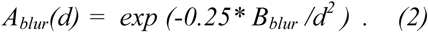

The resolution where the scale factors change from sharpening to blurring is defined by the resolution cutoff (*d*_*cut*_) and the sharpness of the transition is controlled by the transition parameter (*k*). The overall scale factor *A(d)* has the form:

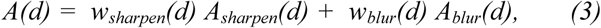

where the resolution-dependent weighting factors *w*_*sharpen*_*(d)* and *w*_*blur*_*(d)* are given by,

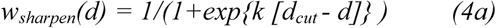

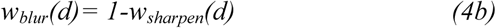

Examining Eq. (4a) it can be appreciated that for low resolution terms where *d* is much larger than *d*_*cut*_, the exponential term will be very small so that the value of *w*_*sharpen*_*(d)* will be close to unity and *w*_*blur*_*(d)* will be close to zero. At the transition resolution *d*_*cut*_, the weights on sharpening and blurring will be equal, and at high resolutions, the blurring weight *w*_*blur*_*(d)* will be close to unity.

Typically the transition resolution *d*_*cut*_ is taken to be the same as the resolution of the map, the blurring B-value has a default value of *B*_*blur*_=200 Å^2^ (a value that leads to blurring by a factor of 250 at a resolution of 3 Å, for example) and the transition parameter has a default value of *k*=10 Å^−1^, leading to a 90% completion of the transition between sharpening and blurring over the resolution range *d*_*cut*_−0.2 Å to *d*_*cut*_+0.2 Å. This leaves the sharpening B-value (*B*_*sharpen*_) as the only parameter that is normally adjusted to optimize a map. Optionally, two of the other parameters can be optimized at present. The keyword “*optimize_d_cut=True*” will optimize the value of *d*_*cut*_, and the keyword “*optimize_k_sharpen=True*” will optimize the value of *k.* These optimizations are off by default because in a large-scale test with 345 cryo-EM maps the average map-model correlation (using B-values of zero for the model) after sharpening with and without these optimizations was essentially identical.

### 2.2. Description of overall map sharpness

We represent a map as a Fourier series and then describe the resolution-dependent increase or decrease in mean amplitudes using an overall “B-factor” (Wlodawer et al., 2008). In essence, this description assumes that the amplitudes of the Fourier terms fall off with resolution according to,

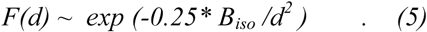

where *F(d)* is the mean amplitude in a shell of resolution *d*, *B*_*iso*_ is the overall B-factor (iso is for isotropic. This B-factor *B*_*iso*_ is calculated for an unsharpened or sharpened map by fitting an anisotropic B-factor to map coefficients (Fourier terms) using the *Phenix* tool *phenix.xtriage* (Zwart et al., 2005) and using the average of the 3 diagonal terms in the matrix representing the anisotropic B-factor as *B*_*iso*_.

### 2.3. Map sharpening by maximization of adjusted surface area

Our core algorithm for map sharpening is to evaluate the interpretability of a sharpened map based on its level of detail and its connectivity (see Fig. 1). The level of detail is derived from the surface area of iso-contour surfaces enclosing a fixed volume of the map (by default 20% of the volume occupied by the macromolecule, see below). The connectivity is derived from the number of contiguous regions enclosed by these iso-contour surfaces. The iso-contour surfaces are simply contours such as those that would be displayed in software such as *Coot* (Emsley et al., 2010) or *Chimera* (Pettersen et al., 2004) at a given threshold level. The surface area (*SA*) of a set of iso-contour surfaces is taken to be the number of grid points that are outside the iso-contour surface and adjacent to a grid point inside the surface. The adjusted surface area (*SA*_*adjusted*_) is the surface area minus the number of contiguous regions (*N*_*regions*_) inside iso-contour surfaces (scaled with a factor *C*_*scale*_ as described below):

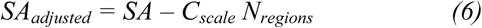

The key parameter in constructing the iso-contour surfaces is the threshold at which the surfaces are drawn. We use a method related to the procedure developed for the *Phenix* tool *phenix.segment_and_split_map* to set this threshold in which the threshold is set to a value that leads to a fixed percentage of the molecular volume inside the contours. To do this requires an estimate of the volume occupied by the molecule(s) in the map. This information can be input directly, but by default we estimate it in two steps. We first assume that some part of the map contains the molecule of interest (and therefore has a high variability of density) and the remainder of the map (often the vast majority of grid points in the map) is essentially empty (flat). The volume occupied by the molecule is identified using tools developed for identification of the region occupied by the macromolecule in crystallographic maps (Wang, 1985, Terwilliger, 1999). The region containing the macromolecule is identified as the region in which the local smoothed squared density is high, indicating a high variability in density. Cryo-EM maps typically have very high contrast between the region containing the macromolecule and the region outside it, so the difference between the regions inside and outside the molecule is generally not difficult to distinguish. The region containing the macromolecule is estimated in a probabilistic fashion based on a guess (in practice, several guesses of widely varying volume fractions) of the volume fraction inside the macromolecule (Terwilliger, 1999). If the volume fraction inside the macromolecule is underestimated, the region identified will typically be larger than the initial estimate. In essence, our procedure consists of identifying the region inside the macromolecule and updating the guess of the volume inside the macromolecule, cycling through this procedure several times. For this calculation, a value of the smoothing radius is needed, and it is chosen by default to be 1.5 times the resolution of the map. (This scale factor for the radius is chosen as a compromise between a high level of detail in the definition of the region inside the macromolecule that would be obtained with a radius equal to the resolution of the map, and a much more robust definition of that region that can be obtained with a larger radius such as twice the resolution of the map.) Once the region containing the macromolecule is identified, the threshold is adjusted to yield a fixed fraction (typically 20%) of the volume of the molecule inside the iso-contour surfaces.

**Figure 1.**
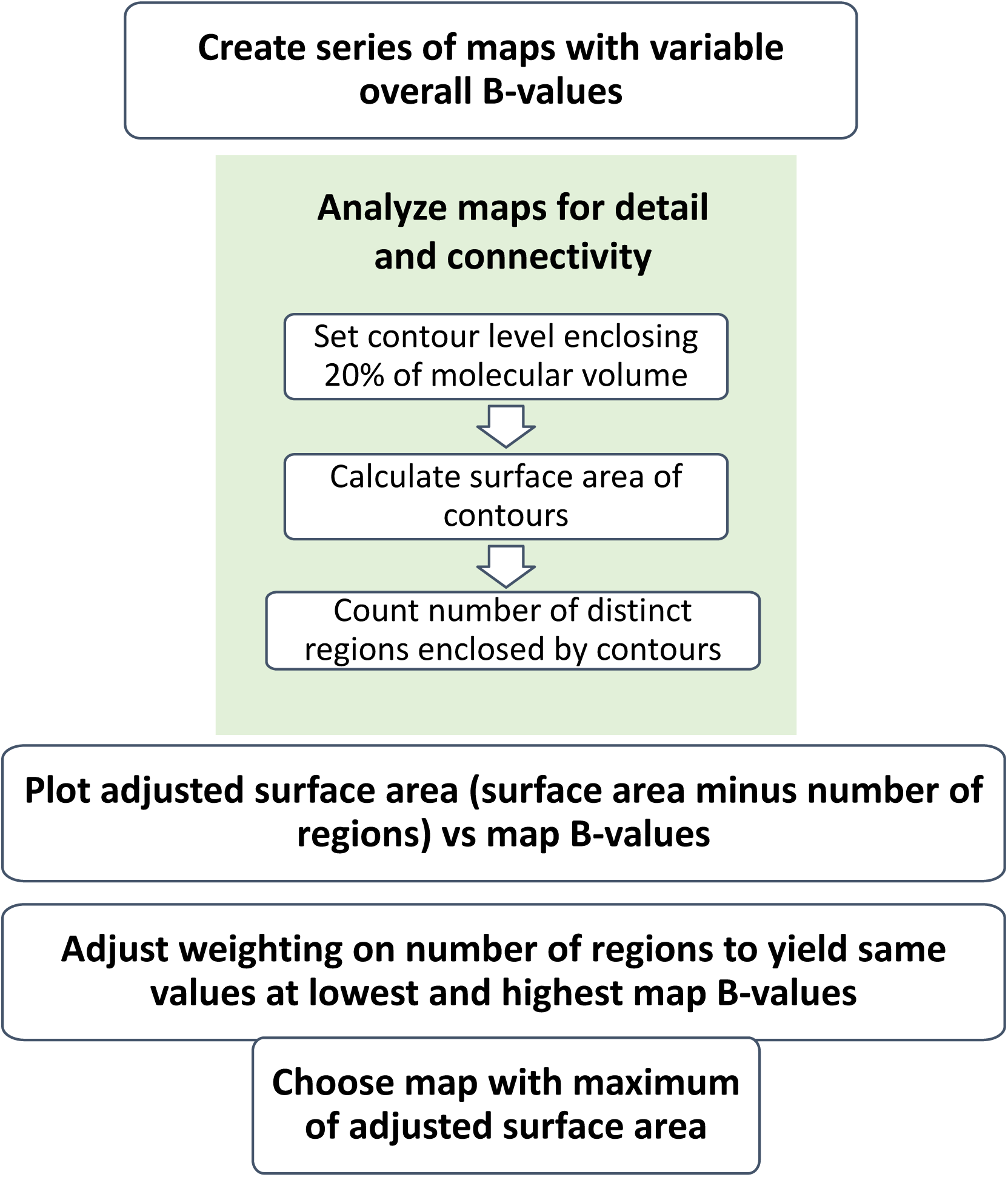
Flow chart for auto-sharpening procedure

### 2.4. Setting the scale factor between surface area and number of regions

In our procedure, the adjusted surface area *SA*_*adjusted*_ is normally calculated over a range of values of sharpening parameters for a particular map. We choose the scale factor *C*_*scale*_ by setting it to a value that yields the same value of the adjusted surface area for the most-sharpened version of a map as the least-sharpened version of the same map. This therefore leads to a set of values of adjusted surface area *SA*_*adjusted*_ *vs* overall B-factor *B*_*iso*_ in which the values of *SA*_*adjusted*_ are the same for the lowest and highest values of *B*_*iso*_. Normally, due to a sharp increase in the number of regions for low values of *B*_*iso*_, the values of adjusted surface area for intermediate values of the overall B-factor are higher than those at the extremes. Our strategy for sharpening is to choose sharpening parameters that maximize the adjusted surface area *SA*_*adjusted*_.

### 2.5. Identifying cases where the algorithm is not applicable

Our procedure of maximizing the adjusted surface area is applicable in cases where the maximum of adjusted surface area occurs at intermediate values of *B*_*iso*_ and where the maximum is clearly identifiable. We identify cases where the approach is not applicable in two ways. First, if the scale factor *C*_*scale*_ would lead to an adjusted surface area at a preset intermediate value of the overall B-factor *B*_*iso*_ (typically *B*_*iso*_ =50 Å^2^) that is less than the values at the extreme values of the overall B-factor *B*_*iso*_, the approach is considered not applicable. Second, we estimate the signal and noise in the adjusted surface area, and we do not apply the procedure if the signal-to-noise is less than a preset ratio (typically 3:1). The signal is taken to be the difference between the maximum value of the adjusted surface area and the value of the adjusted surface area at the extremes of *B*_*iso*_. The noise is estimated from the local variation of adjusted surface area for adjacent values of *B*_*iso*_. The local variation of adjusted surface area is taken to be the difference between the value of adjusted surface area *SA*_*adjusted*_ at a particular value *B*_*iso*_ and the value of *SA*_*adjusted*_ interpolated between the neighboring two values of *B*_*iso*_. The noise is then taken to be the rms value of this local variation of adjusted surface area.

### 2.6. Map sharpening based on kurtosis, half-map correlation, or a model

We implemented methods for map sharpening based on kurtosis, the correlation between half-maps, and on map-model correlation. For sharpening based on kurtosis the overall sharpening B-value was simply adjusted to maximize the kurtosis of the map.

For half-map and map-model correlation a procedure similar to that developed by Rosenthal and Henderson (2003) was applied, except that the resolution dependence of a model-based map was used as a normalization factor in each case (this normalization factor is similar to a scale factor used by Jakobi et al., 2017). The map-map or map-model Fourier Shell Correlation (FSC) was calculated in shells of resolution. For half-map correlations, this correlation (CC) was converted into an estimate of the true correlation CC^*^ between the full map and a perfect map using the formula (Rosenthal and Henderson, 2003),

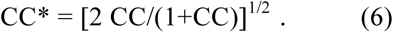

For map-model correlations, by analogy to the σ_A_ analysis of errors in a crystallographic map (Read, 1986), the correlation (CC) was assumed to be related to the true correlation CC^*^ by an exponential error function,

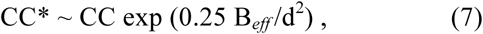

where the effective B-value (B_*eff*_) is estimated from a guess of the error in the model (rms_e_, assumed to be ¼ the resolution of the map) using the relation,

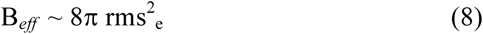

Finally, for each shell of resolution, the ratio (*R*) of the mean amplitude in the Fourier coefficients for a model-based map calculated with B-values of zero to the mean amplitudes of the Fourier terms representing the full starting map is calculated. Then for half-map and map-model sharpening, the scale factor applied to all amplitudes in this shell of resolution is R CC^*^. That is, amplitudes are increased for shells where the model-based amplitudes calculated with a B-value of zero are larger than those obtained from the original map, and decreased for shells where the estimated accuracy of the map is low.

### 2.7. Local map sharpening

We applied sharpening to overlapping local regions in a map and combined the resulting partial maps to yield a locally-sharpened map. In order to focus on regions containing density, the original map was first analysed with the *Phenix* tool *phenix.segment_and_split_map* to obtain a set of regions of density representing most of the density in the map. Then an overlapping set of boxes, covering these regions of density, were extracted from the original map. The size of these boxes was typically 40 grid units on a side. The map corresponding to each box was sharpened, yielding a set of overlapping sharpened maps. The maps were combined by weighting overlapping regions based on the distance of each grid point from the center of the corresponding box, with an exponential fall-off of the weighting factor *w*,

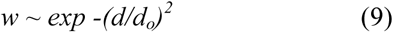

where the characteristic distance *d*_*o*_ is by default set to be the average nearest-neighbor distance between centers of boxes.

### 2.8. Map-model correlation using a model with B-values of zero as a metric of map quality

It is useful to have a metric of map quality in order to compare different approaches for map sharpening. For this work the metric of map quality is the correlation between the map of interest and a map calculated from a model in which all the B-factors are set to zero, and using data to the same resolution as used in the map of interest. (This is done by taking the original map, converting it into a box of Fourier map coefficients, removing all terms that have resolution higher than the defined value, and finally computing a real-space map from the truncated set of Fourier map coefficients.) The calculation of this map-model correlation is carried out using the *Phenix* tool *phenix.map_model_cc* using a model that has B-factors set to zero (for example using the *Phenix* tool *phenix.pdbtools* with the keyword *set_b_iso=0*) and specifying the same resolution as is used in calculation of the map of interest.

### 2.9. Map-model pairs chosen from the EM Data Bank and the Protein Data Bank

We began with all 1097 maps in the EMDB as of August, 2017 that had an associated model in the PDB. We then removed 91 map-model pairs for which the resolution reported in the PDB and the EMDB differed by 0.2 Å or more or was not reported, and selected all the remaining pairs for which the reconstruction resolution was 4.5 Å or better, yielding 401 map-model pairs. We then removed 24 additional map-model pairs for which the map-model correlation was less than 0.3. Finally, we removed 16 map-model pairs for which the signal-to-noise criterion for applying our procedure was not satisfied (see §2.5), leaving 361 map-model pairs at resolutions from 1.8 Å to 4.5 Å. For some analyses only a subset of these datasets were tested (the comparison of automatic sharpening with maximization of model-map correlation used 345 of the datasets and comparisons involving half-dataset correlations only include the 59 datasets in our sample that had half-datasets deposited).

For each map-model pair, the region of the map near the model was extracted with the *Phenix* tool *phenix.map_box*, including density within 5 Å of any atom in the model, and placed in a new (typically smaller) box with the same gridding as the original map, and a new origin at one corner of the map. This map and associated model (translated as necessary to match the new map) were used in the analyses described here.

### 2.10. Software availability

The automatic sharpening tool *phenix.auto_sharpen* is available as part of the *Phenix* software suite.

## 3. Results and Discussion

### 3.1. Optimizing parameters for map sharpening/blurring by maximization of adjusted surface area

The key idea in this work is that a map that is optimally sharpened will have more detail, leading to a high surface area of an iso-contour surface, yet at the same time it will have a high degree of connectivity, leading to a low number of regions enclosed by that same iso-contour surface. Figs. 2 and 3 illustrate these relationships for the cryo-EM map of the anthrax protective antigen pore (Jiang et al., 2015) with a map (emd_6224) deposited in the EM Data Bank (EMDB, Lawson et al., 2016) and a model (3j9c) deposited in the Protein Data Bank (PDB, Berman et al., 2000; Bernstein et al., 1977) and an analysis at a nominal resolution of 2.9 Å. In each panel of Fig. 2 the deposited map is represented as a Fourier series up to a resolution of 2.9 Å, and the amplitudes are adjusted with various “sharpening” B-factors that emphasize or deemphasize high-resolution information in the map.

**Figure 2.**
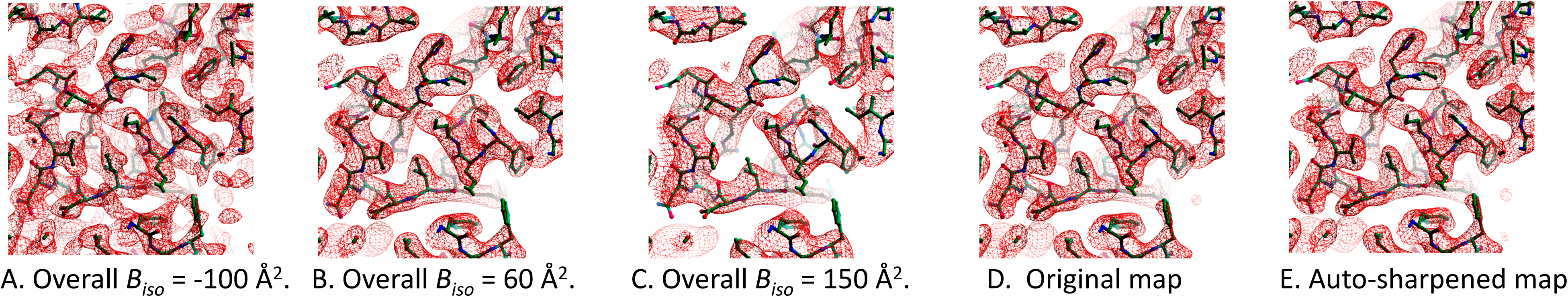
Sharpened/blurred versions of cryo-EM map of the anthrax protective antigen pore (Jiang et al., 2015). A. Overall *B*_*iso*_ = −100 Å^2^. B. Overall *B*_*iso*_ = 60 Å^2^. C. Overall *B*_*iso*_ = 150 Å^2^. D. Original map (emd_6224). E. Auto-sharpened map, overall *B*_*iso*_ = 20 Å^2^. All contours drawn at 3σ. Maps drawn with *Coot* (Emsley et al., 2010).

**Figure 3.**
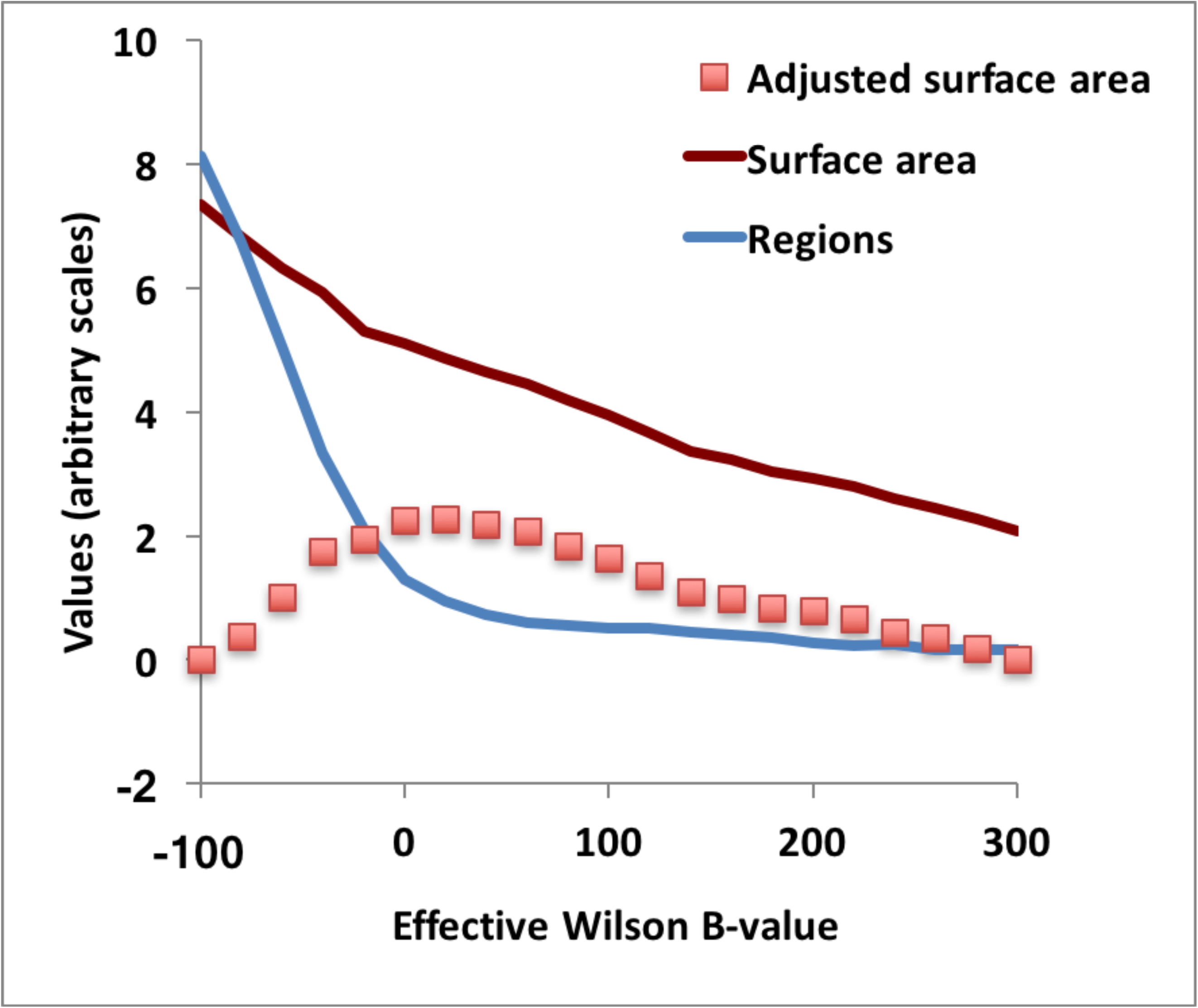
Surface area, number of regions, and adjusted surface area of sharpened/blurred versions of cryo-EM map of the anthrax protective antigen pore (Jiang et al., 2015). See text for details.

In Fig. 2A, the map is sharpened so that the overall resolution dependence is approximated by a B-factor of −100 Å^2^ (the amplitudes strongly increase at high resolution). In Fig 2B, the B-factor is 60 Å^2^ (the amplitudes fall off somewhat high resolution) and in Fig. 2C, it is +150 Å^2^ (the amplitudes fall off substantially at high resolution). Examining Fig. 2A, it can be seen that this highly sharpened map has a high level of detail but also is somewhat fragmented (the map has some breaks where there is no density in the locations of atoms in the model). Framed in terms of adjusted surface area, this map has a high surface area of the iso-contour surfaces, but the map also has a large number of separate regions that are enclosed by these surfaces. Fig. 2B still has a high level of detail but has less fragmentation. Fig. 2C has much less detail and correspondingly less surface area of the iso-contour surfaces, but it also has less fragmentation than either Figs 2A or 2B.

Fig. 3 quantifies this analysis of maps in Fig 2. Fig. 3 shows the surface area, number of regions, and adjusted surface area for a series of such maps with overall B-factors varying from −100 Å^2^ to 300 Å^2^. It can be seen that as the overall B-factor for the map decreases (leading to an emphasis of high-resolution information), the surface area of the map increases. The number of regions inside iso-contour surfaces increases in general as well, and in particular the number of regions become much larger very rapidly for maps with very low (or negative) overall B-values. This rapid increase in the number of regions can be understood in terms of over-sharpening of the map and the associated fragmentation of regions of density.

The adjusted surface area is the surface area less a constant times the number of regions. This constant is set (see Methods) for a particular map to yield the same value of the adjusted surface area for sharpening with the extreme low and high B-values considered. It can be seen that after setting this constant (*C*_*scale*_), the adjusted surface area *SA*_*adjusted*_ increases as a function of the overall B-value of the map, comes to a maximal value, and then decreases. In our procedure, the optimal value of sharpening parameters is the value that leads to a maximum of the adjusted surface area *SA*_*adjusted*_. For the map of the anthrax protective antigen pore, this leads to an optimized map with an overall B-value of about 20 Å^2^ (Fig. 2E).

Fig. 4 illustrates the resolution dependence of the mean amplitude of Fourier coefficients for the deposited map of the anthrax protective antigen pore and compares them with those of the auto-sharpened map obtained on the basis of Fig. 3. It can be seen that in this case the auto-sharpened map has a resolution dependence of amplitudes that is similar to that of the deposited map, except that due to the blurring of the map near the high-resolution limit the amplitudes for the auto-sharpened map fall off in this resolution range. Figs. 2D and 2E show that the overall appearance of the deposited and auto-sharpened maps are quite similar in this case.

**Figure 4.**
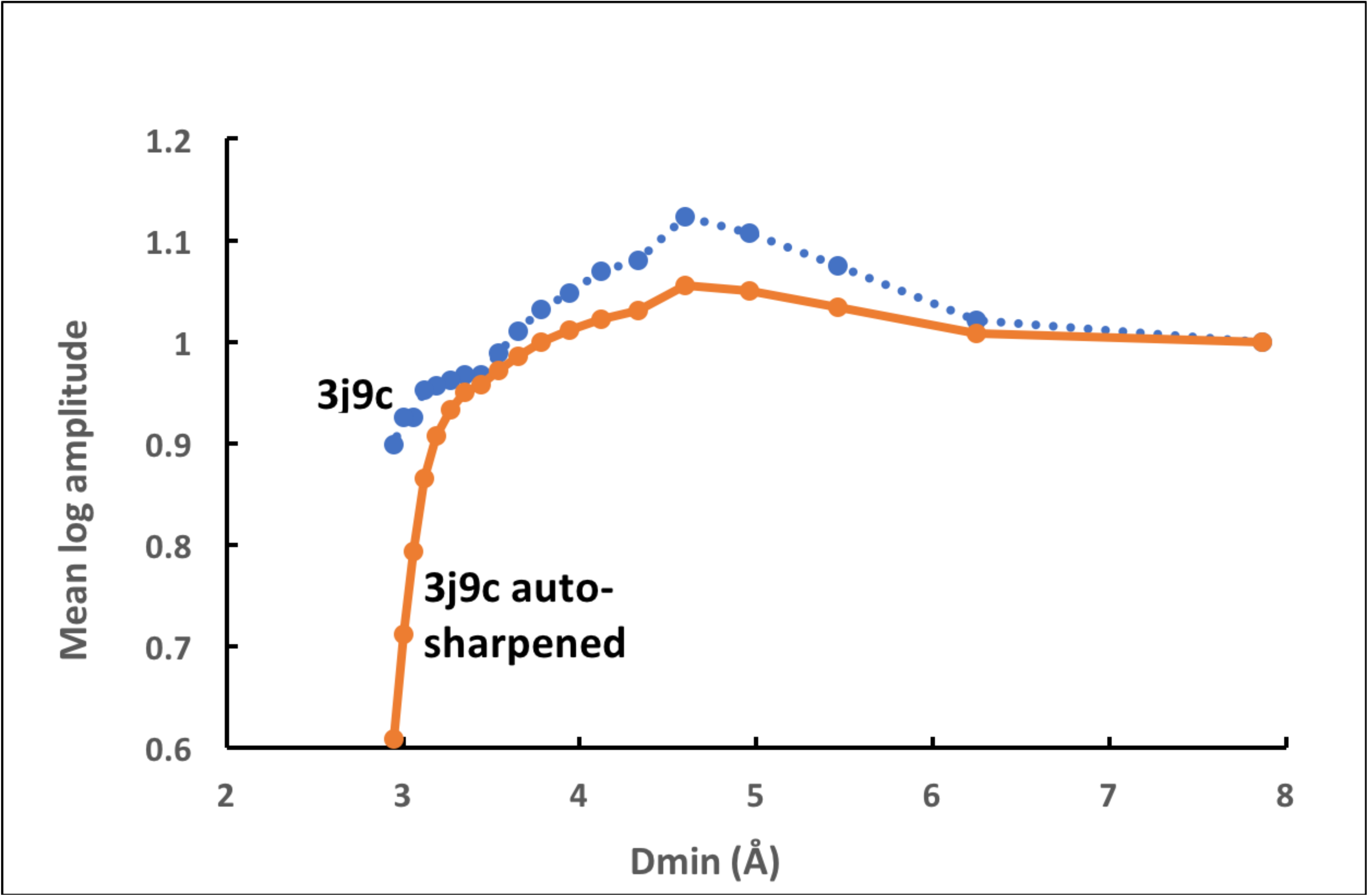
Resolution dependence of the mean amplitude of Fourier coefficients for cryo-EM map of the anthrax protective antigen pore and for auto-sharpened map.

### 3.2. Map-model correlation using a model with B-factors of zero as a metric of map quality

It would be useful to have an overall metric of map quality that could be used to evaluate the quality of maps sharpened with our automated methods. As the maps we are analyzing in our tests have already been interpreted, structural models are available and can be used for this purpose. A challenge in map evaluation, even having a structural model, is that it is not exactly clear on what basis the map should be evaluated. Presumably the map should be evaluated based on its interpretability, but this could be visual interpretability, or the quality of model that can be automatically built into the map, or some other measure of interpretability.

For crystallographic maps, a commonly-used metric of map quality is simply the map-model correlation (e.g., Afonine et al., 2018). This is calculated by creating a model-based map and obtaining the correlation between density values in the model-based map and those in the map of interest (see Urzhumtsev et al., 2014 for an analysis of related metrics). Such a map-model correlation may be calculated for the map as a whole or just for grid points near atoms in the model. Normally the atomic model used in this calculation will have been refined against X-ray data and will therefore have refined values of the B-factors that match the resolution-dependence of the measured X-ray (intensity) data. The B-factors used to generate the model-based map are normally the B-factors that are associated with the refined model, and in this way the resolution dependencies of the model-based map and of the original data are similar.

For cryo-EM maps the situation is more complicated because atomic models are typically refined, not against primary data, but rather by comparison with a map that been modified with resolution-dependent scaling factors (e.g., Rosenthal and Henderson, 2003). Additionally, deposited maps may or may not be the maps used in refinement or map interpretation. Consequently, the B-factors in refined cryo-EM maps do not necessarily represent the resolution dependence of the original images used in the reconstruction or that of the deposited maps.

To sidestep this issue, in this work we calculate model-based maps using B-factors of zero. This map of course does not depend on the values of the B-factors in the model and therefore it avoids the problems associated with values of refined B-factors. Further, it has the advantage that if the model were perfect, this model-based map would (more or less) be the clearest map that could be created to represent it. The quality of a particular map of interest is then evaluated by calculating its correlation with such a zero B-factor model-based map, considering only points in the map that are near atoms in the model. (An alternative approach to using a model to evaluate a map would be to adjust the B-factors in the model to maximize the map-model correlation. It is important however to note that in the present work the goal is to identify the optimal sharpening of a map, not to identify whether the map is accurate. In this context, the approach of adjusting B-factors in the model to maximize the map-model correlation has the disadvantage that a map with slightly more accurate low-resolution information than high-resolution information could have a maximal map-model correlation when it is highly blurred, whereas the most useful map would be one that includes the high-resolution information.)

The key question about using model-map correlations based on a model with zero B-factors to evaluate maps is whether this metric can in fact identify maps that have high interpretability. We attempt to address this issue here using two sets of variably-sharpened maps. We first examine the maps visually, evaluating them based on connectivity of density along the polypeptide chains and on the level of detail in the density. Then we evaluate them based on map-model correlations. Figs. 5 and 6 each show a series of maps that has been sharpened using Eq. 3 with a variety of sharpening B-values, leading to overall B-values of the maps that range from 0 Å^2^ to 120 Å^2^. Examining the series of maps with variable sharpening in Fig. 5 (innexin-6 gap junction channel; emd_9571 and PDB entry 5h1r; Oshima et al., 2016), it can be appreciated that the map in Fig. 5E (or perhaps Fig. 5D) is probably the easiest to interpret, in that the density most closely matches the model and there are no breaks in the density along the polypeptide chain. These maps have overall B-values of 80 Å^2^ and 60 Å^2^, respectively. For Fig. 6 (TRPV1 channel; emd_5778 and PDB entry 3j5p; Liao et al., 2013) the situation is somewhat different, and for this map the same criteria plausibly lead to the conclusion that the map in Fig. 6B, with an overall B-value of about 20 Å^2^, is most useful.

**Figure 5.**
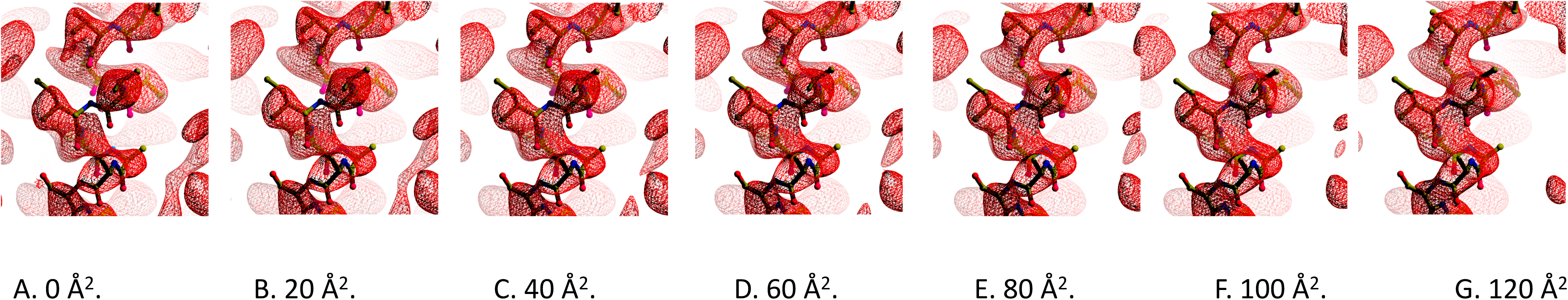
Cryo-EM map of the innexin-6 gap junction channel (emd_9571 and PDB entry 5h1r; Oshima et al., 2016) after application of sharpening. Contours are chosen to yield equal volume inside contours in each map (Urtzhumtsev et al. 2014). Final values of overall *B*_*iso*_ and contour levels used are: A. 0 Å^2^, 1.20 σ. B. 20 Å^2^, 1.20 σ. C. 40 Å^2^, 1.20 σ. D. 60 Å^2^, 1.22 σ. E. 80 Å^2^, 1.23 σ. F. 100 Å^2^, 1.24 σ. G. 120 Å^2^, 1.26 σ. Maps drawn with *Coot* (Emsley et al., 2010) and *Raster3D* (Merritt and Bacon, 1997).

**Figure 6.**
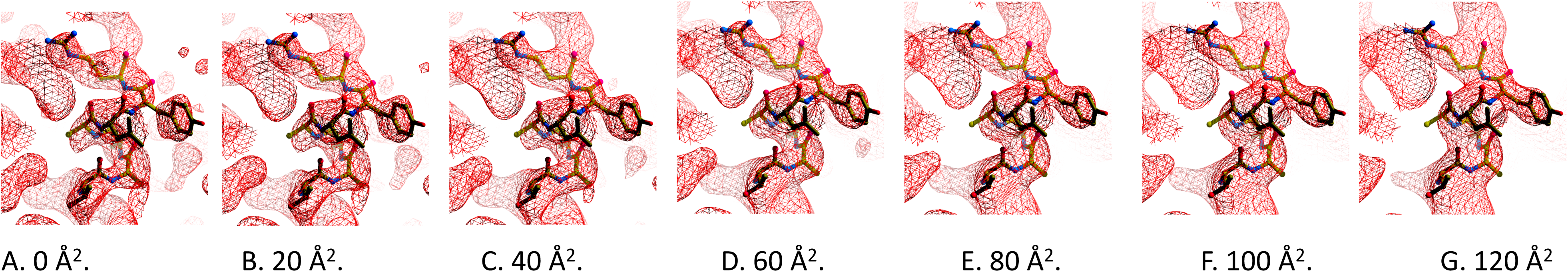
Cryo-EM map of the TRPV1 channel (emd_5778 and PDB entry 3j5p; Liao et al., 2013) in the vicinity of residues 449-455 after application of sharpening. Contours are chosen to yield equal volume inside contours in each map (Urtzhumtsev et al. 2014). Final values of overall *B*_*iso*_ and contour levels used are: A. 0 Å^2^, 1.19 σ. B. 20 Å^2^, 1.20 σ. C. 40 Å^2^, 1.24 σ. D. 60 Å^2^, 1.30 σ. E. 80 Å^2^, 1.36 σ. F. 100 Å^2^, 1.41 σ. G. 120 Å^2^, 1.49 σ. Maps drawn with *Coot* (Emsley et al., 2010) and *Raster3D* (Merritt and Bacon, 1997).

We then evaluated each map in Figs. 5 and 6 based on map-model correlation. Fig. 7A shows an analysis of the maps in Fig 5 in which the map-model correlation is plotted as a function of the overall B-value for sharpened maps. Consider first the points labelled “*model b_iso 0*”. These reflect the map-model correlation as a function of overall map B-values, where the model map is calculated with B-values (atomic displacement factors) of zero. It can be seen that the map-model correlation has a maximum at an overall map B-value of 90-100 Å^2^, similar to the visually optimal overall map B-value of about 80 Å^2^. Next note that if the same analysis is done using model-map correlations calculated with model B-values of −100 Å^2^, the map-model correlation has a maximum at an overall map B-value of 40-50 Å^2^, and if the model B-values are set to +100 Å^2^, the map-model correlation has a maximum at an overall map B-value of 160 Å^2^. This case illustrates that the values used for the model B-values have a very large effect on the overall map B-value that leads to a maximal model-map correlation. It also shows that using model B-values of zero lead to a maximal model-map correlation at overall map B-values that are similar to those identified as optimal by visual inspection.

**Figure 7.**
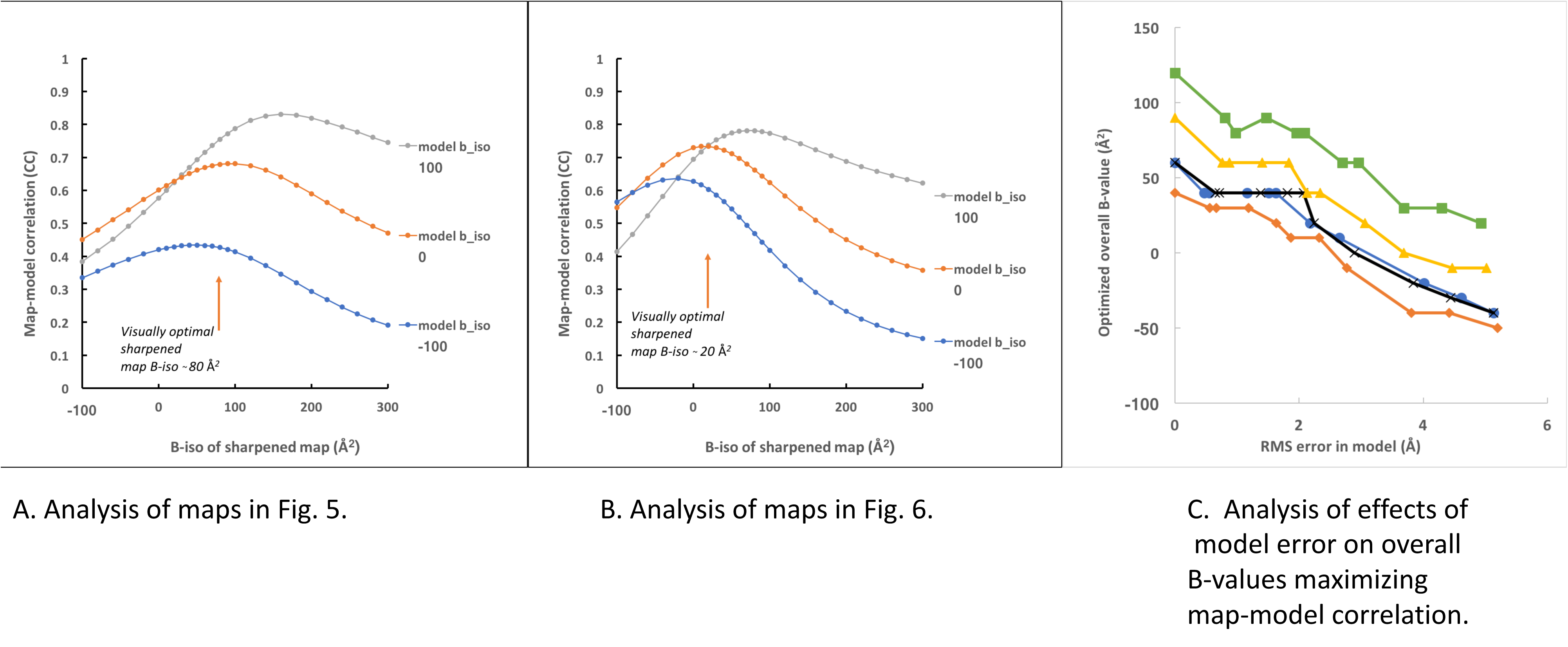
Map-model correlation as a function of overall B-value for sharpened maps. A. Analysis of maps in Fig. 5. B. Analysis of maps in Fig. 6. C. Analysis of effects of model error on overall B-values maximizing map-model correlation. See text for details.

In Fig. 7B we apply the same analysis to the maps from Fig. 6. The model B-values of zero lead to a maximal model-map correlation at an overall map B-value (*B*_*iso*_=10-20 Å^2^) that is similar to the value identified as optimal by visual inspection (*B*_*iso*_=20 Å^2^). These results suggest that map-model correlation calculating the model-based map using model B-values of zero can reproduce visual evaluations of map interpretability quite well.

### 3.3. Model accuracy required for evaluating maps based on map-model correlation with zero B-factors

An important caveat in using map-model correlation with model B-values of zero as a metric of map interpretability is that the model has to be similar to the true structure. We carried out a simulation to get an idea of how accurate the model has to be in order for these map-model correlations to identify interpretable maps. We took chain A of the major capsid protein of rotavirus (PDB entry 1qhd, Mathieu et al., 2001) and used it to calculate model-based Fourier coefficients to a resolution of 3.5 Å using an overall B-value of 72 Å^2^. Then we randomized the phases, introducing variable resolution-dependent errors using *RESOLVE* (Terwilliger, 1999) to yield a series of maps of varying quality. We then began with each map of variable quality and sharpened or blurred the map to yield modified maps with a range of overall B-values. We began again with the same maps of variable quality, and for each we generated a series of new models by randomized the starting model with the *shake* algorithm in the *Phenix* tool *phenix.pdbtools*, refining it against the map with the *Phenix* tool *phenix.real_space_refine*. This process yielded (1) a set of maps with variable quality, and for each such map, (2) a set of sharpened/blurred maps with varying overall B-values, and (3) a set of refined models with varying coordinate errors.

We then used these maps and models to determine how accurate a model had to be for the maximum of map-model correlation occurred at a very different overall map B-value than the value obtained when there was no error in the model. We started with the original model-based map without errors along with its corresponding sharpened/blurred versions with varying overall B-values and its corresponding refined models with varying coordinate errors. For each model with varying coordinate errors, we set the model B-values to zero and carried out an analysis similar to the one used in Figs. 7A and 7B, yielding a plot of map-model correlation as a function of overall map B-value and a value of the overall map B-value (*B*_*iso*_) that maximized map-model correlation. Fig. 7C (blue curve) shows these overall map B-values maximizing map-model correlation as a function of the rms error in the model coordinates. It can be seen that the optimal map B-value is about 60 Å^2^ for the perfect map and perfect model. For models with increasing error, the overall B-value maximizing map-model correlation gradually decreases, changing by about −20 Å^2^ from its initial value when the rms error increases to about 1 Å and by −60 Å^2^ when the rms error increases to 2 Å.

We repeated this analysis using maps of variable quality, with map-model correlation (CC) between the original model with B-values set to zero and the maps of variable quality ranging between 0.42 and 0.98 (Fig. 7C; blue, CC=0.98; orange, CC=0.78; black, CC=0.63; yellow, CC=0.50; and green, CC=0.42). The optimal overall B-values for a perfect model (ordinate of zero in Fig. 7C) gradually increase with decreasing map quality, as expected. The overall B-values maximizing map-model correlation gradually decrease with increasing model error in much the same way that they did for the perfect map. As maps that differ in overall B-value by 20 Å^2^ are very similar and even those differing by 60 Å^2^ are not greatly different (see Figs. 5 and 6), this analysis suggests that models that have coordinate errors of up to about 1.5-2.0 Å, or about half the resolution of this 3.5 Å map, could be used effectively for evaluating the quality of this map.

### 3.4. Evaluation of adjusted surface area maximization as a method for optimizing map interpretability

We applied our algorithm for automatic map sharpening to 345 of the map-model pairs from the EMDB and PDB identified above. We then compared the overall B-values obtained using our procedure with those obtained by maximizing map correlation to a model-based map calculated with model B-values of zero (Fig. 8). It can be seen that the automatic map sharpening procedure yields overall B-values similar to those obtained by maximizing map-model correlation. The rms difference between overall B-values using the two approaches is 32 Å^2^, which by reference to Figs 5 and 6, would lead to relatively small differences in map appearance.

**Figure 8.**
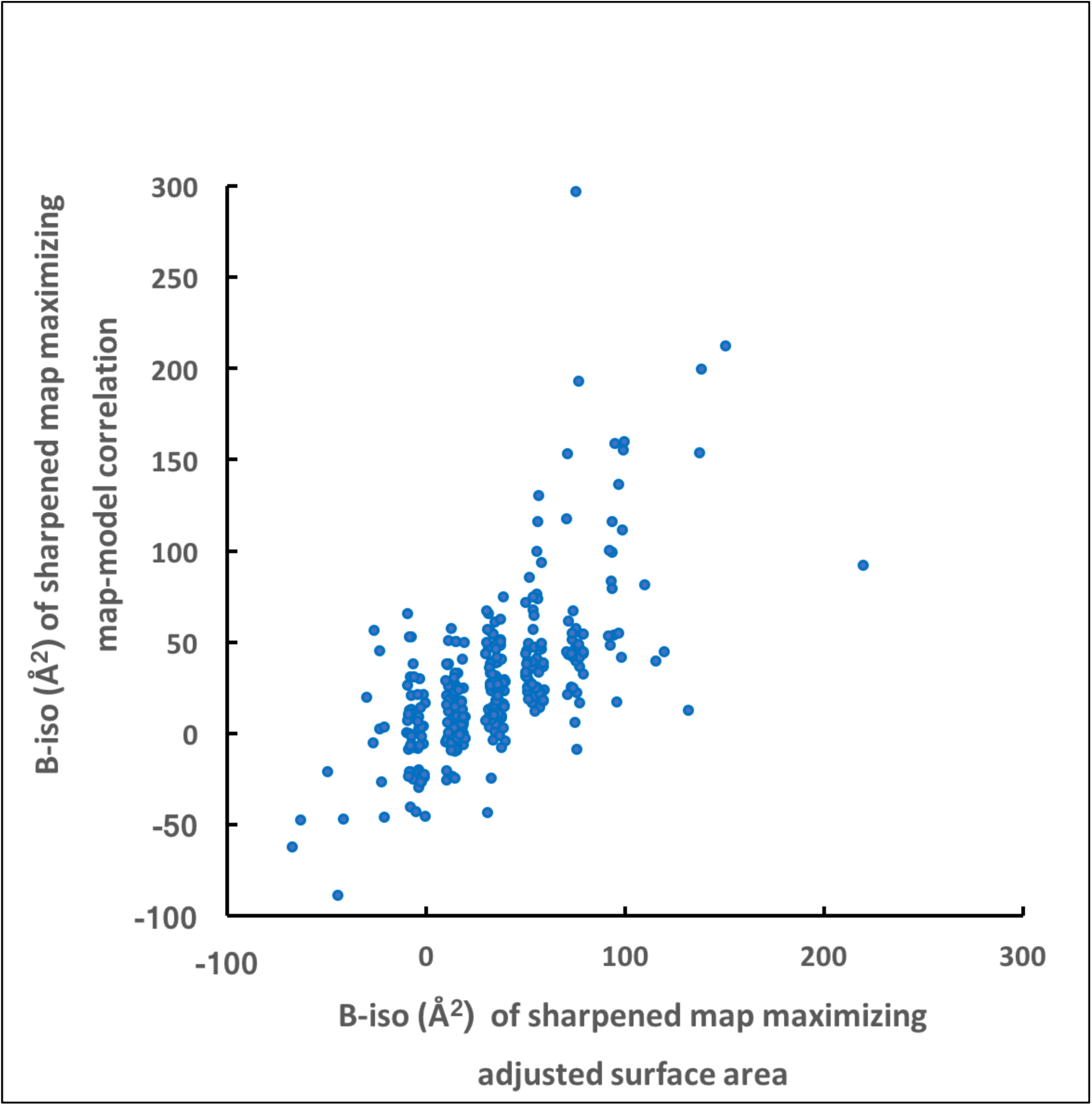
Comparison of overall B-values obtained by automatic sharpening with those obtained by direct maximization of map-model correlation.

We next compared maps obtained using our automatic map sharpening procedure with the maps deposited in the EMDB. In each case we used the map-model correlation calculated with model B-values of zero as our metric of map quality. Fig. 9A compares our adjusted surface area-based maps for 361 deposited map-model combinations with the original maps. It can be seen that in a high fraction of cases (65%) the automatically sharpened maps have a map-model correlation that is better than that of the deposited maps, only 4% have a correlation more than 0.02 worse than the deposited maps while 28% have a correlation at least 0.02 higher. Overall, the mean correlation is 0.02 higher for automatic sharpening. Considering that the deposited maps are generally already optimized by the depositors, this indicates that the automatic sharpening procedure works well.

**Figure 9.**
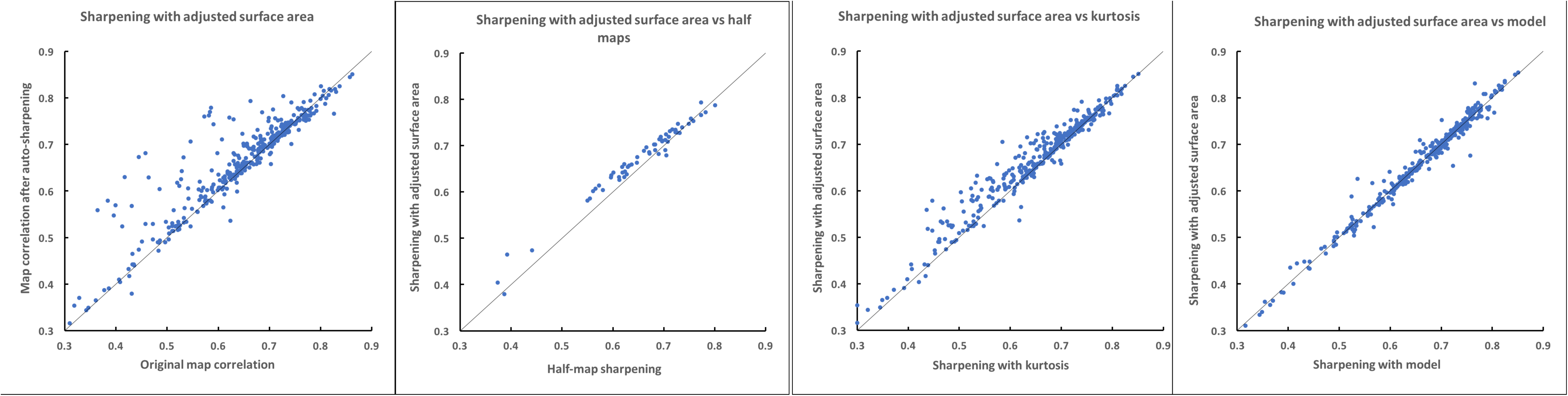
Map-model correlation for automatically-sharpened and deposited cryo-EM maps. A. Comparison of original maps and maps sharpened by maximization of adjusted surface area. The ordinate is the map-model correlation (calculated using model B-values of zero) for the deposited maps; the abscissa is the map-model correlation for the automatically sharpened maps. B. Comparison of maps sharpened using half-map based sharpening with maps sharpened with maximization of adjusted surface area. C. Comparison of maps sharpened using maximization of kurtosis with maps sharpened with maximization of adjusted surface area. D. Comparison of maps sharpened using of map-model correlation with maps sharpened with maximization of adjusted surface area.

### 3.5. Comparison of methods for optimizing map interpretability

We compared our procedure for map sharpening based on adjusted surface area with other procedures for map sharpening. Fig. 9B illustrates that map sharpening using our adjusted surface area approach yields higher map-model correlation (0.02 better on average) compared to using half-map based sharpening (Rosenthal and Henderson (2003). Fig. 9C illustrates that map sharpening using our adjusted surface area approach also yields higher map-model correlation (0.01 better on average) compared to using a target of the kurtosis of the map. Fig 9D shows that model-based optimization is just about the same as (no difference in average map-model correlation) our adjusted surface area-based procedure.

### 3.6. Local sharpening of maps

Jakobi et al. (2017) have recently described a method for local sharpening of maps that uses a refined model and is based on using the local resolution-dependence of the model-based map to normalize Fourier terms in a map of interest. They find that some regions of maps can be more interpretable when local sharpening is applied than when global sharpening is used. We have implemented a local map sharpening algorithm and it can be applied along with any of our approaches for map sharpening. In our implementation of local sharpening we have not seen any substantial differences compared with global sharpening. Fig. 10A compares global and local sharpening using maximization of adjusted surface area. Fig 10B compares them using half-map sharpening, and Fig. 10C compares them with model-based sharpening. For maximization of adjusted surface area and model-based sharpening the maps have very similar overall correlations to model-based maps calculated with B-values of zero, indicating that at least overall, the maps are about equally useful. For half-map sharpening the locally-sharpened maps are slightly better on average than the globally sharpened maps (these locally-sharpened maps are still not quite as good as those obtained by maximization of adjusted surface area, however). As our results are specific to the methods for local sharpening described here, it is entirely possible that a different implementation might yield more substantial improvements in map quality.

**Figure 10.**
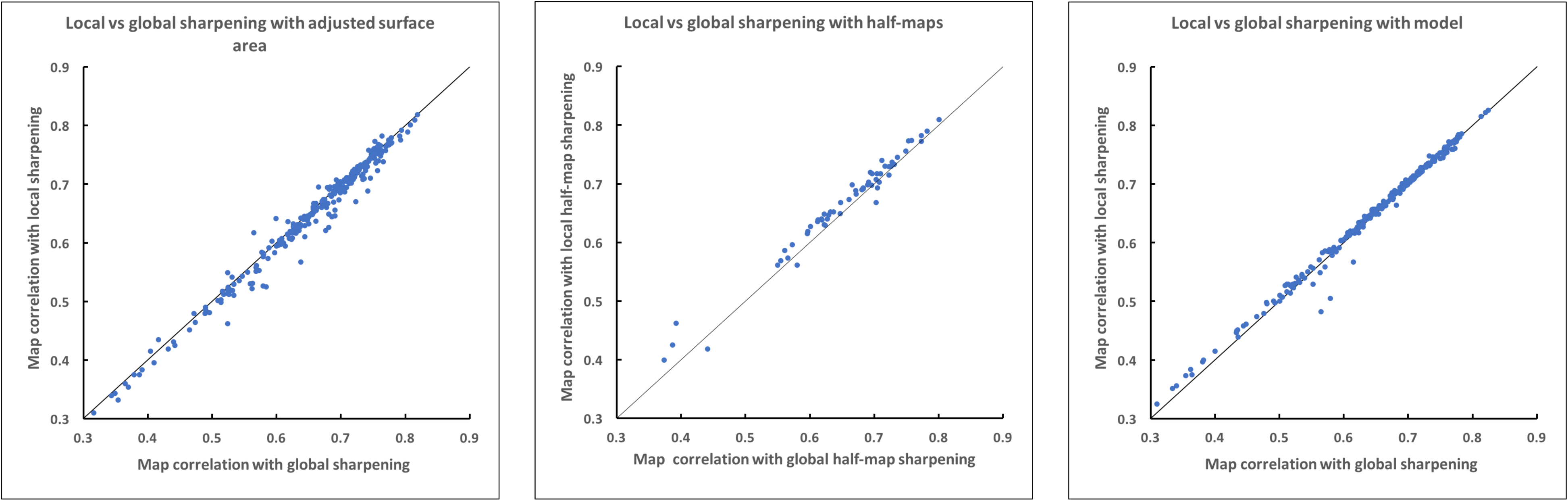
Comparison of map-model correlations using global vs local sharpening. A. Sharpening using maximization of adjusted surface area. B. Sharpening using half-map correlation. C. Model-based sharpening.

### 3.7. Examples of map improvement with auto-sharpening

Fig. 11 illustrates the map improvement that can be obtained with auto-sharpening. The purpose of this Figure is to show that the auto-sharpening procedure can automatically provide highly interpretable maps without manual intervention. The examples illustrated were chosen based on the increase in map-model correlation after sharpening, so they reflect the maximum improvement that might be anticipated, and similar results could presumably be obtained by careful manual adjustment of sharpening parameters as well. For each case, Fig. 11 shows a section of the original map (e.g., Fig. 11A) and the same section of the auto-sharpened map (e.g., Fig. 11B), along with the overall B-values associated with each map and the contour level used to illustrate the map. It can be seen that the improvement in map interpretability through automatic sharpening can be considerable.

**Figure 11.**
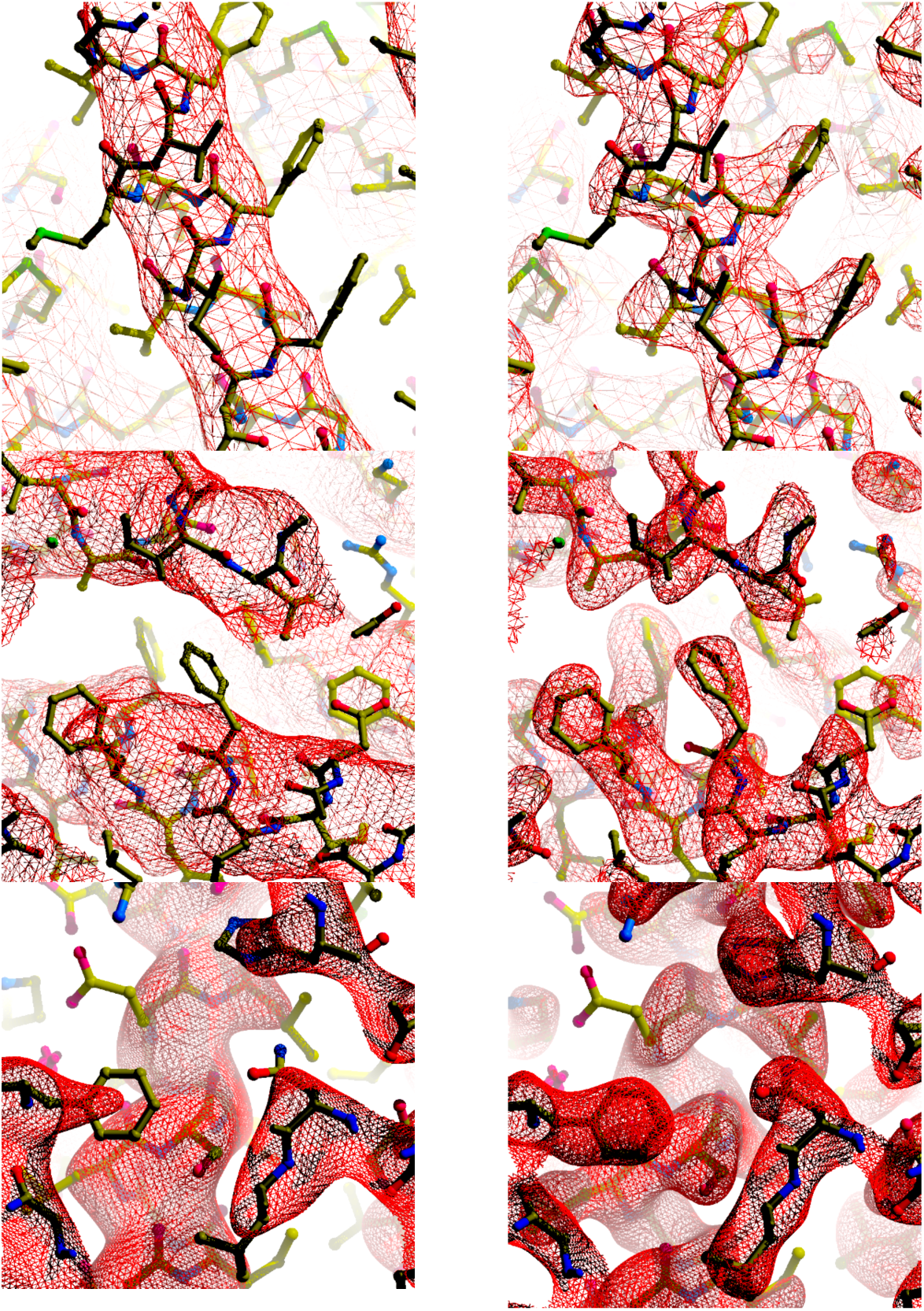
Auto-sharpening applied to maps from EMDB. Maps shown are among those that showed the most improvement in map correlation using *phenix.auto_sharpen*. Contours are chosen to yield equal volume inside contours in each map (Urtzhumtsev et al. 2014). Maps drawn with *Coot* (Emsley et al., 2010) and *Raster3D* (Merritt and Bacon, 1997). A, B. Map of the high-conductance Ca(2+)-activated K(+) channel (emd_8414 and PDB entry 5tji; Hite et al., 2017) before and after application of sharpening. Original and final values of overall *B*_*iso*_ and contour levels used are: A. 260 Å^2^, 1.57 σ. B. 20 Å^2^, 1.5 σ. C, D. Map of the cystic fibrosis transmembrane conductance regulator (emd_8461 and PDB entry 5uar; Zhang and Chen, 2016) before and after application of sharpening. Original and final values of overall *B*_*iso*_ and contour levels used are: C. 290 Å^2^, 1.01 σ. D. −60 Å^2^, 1.0 σ. E, F. Map of lactate dehydrogenase (emd_8191 and PDB entry 5k0z; Merk et al., 2016) before and after application of sharpening. Original and final values of overall *B*_*iso*_ and contour levels used are: E. 102 Å^2^, 2.47 σ. F. −20 Å^2^, 2.0 σ.

### 3.8. Conclusions

Our major observation is that it is possible to automatically identify optimized sharpening parameters for cryo-EM maps by simultaneously maximizing the level of detail in the maps and the connectivity of the maps. The adjusted surface area of a map reflects these factors, does not require a model or any other prior interpretation of a map, and maximizing it leads to improved maps. A secondary observation is that a useful metric of cryo-EM map quality is the correlation between the map and a model-based map in which the atomic B-values all have values of zero. Applying our automatic map sharpening procedure to 361 cryo-EM maps with resolutions from 1.8 Å to 4.5 Å and evaluating the resulting maps with this metric, we find that our procedure can improve deposited maps, that it is an improvement over both kurtosis-based and half-map sharpening procedures, and that it is about equal overall to model-based map sharpening, all as implemented in the *Phenix* tool *phenix.auto_sharpen.* The focus in this work is on optimization of cryo-EM maps, but the *phenix.auto_sharpen* tool can be applied as well to X-ray crystallographic maps. We have carried out limited tests on crystallographic cases but it seems likely that the approaches will generally be suited to both cryo-EM and crystallographic maps. Two important differences, however, are that for crystallographic maps in the early stages of structure determination the maps may be very poor and the region occupied by the macromolecule may be very indistinct. This could lead to poorer identification of the region where the macromolecule is located and less distinction between maps that are more or less interpretable as each may be highly fragmented. Nevertheless, the auto-sharpening of maps may be a useful addition to automated structure determination tools for X-ray crystallography. As noted in the Introduction, the *phenix.autosol* tool for automated structure determination (Terwilliger et al., 2009) by experimental phasing and the *phenix.autobuild* tool for iterative model-building, density modification and refinement (Terwilliger et al., 2008), the current default is to sharpen maps using an empirical formula based on the resolution of the map. Applying an optimization technique such as the one developed here could potentially improve both of these steps in structure determination.

## Acknowledgements

The authors appreciate support received from the US National Institutes of Health (grant P01GM063210 to P.D.A. and T.C.T.). This work was partially supported by the US Department of Energy under contract DE-AC02-05CH11231.

